# RAB18 impacts autophagy via lipid droplet-derived lipid transfer and is rescued by ATG9A

**DOI:** 10.1101/421677

**Authors:** Fazilet Bekbulat, Daniel Schmitt, Anne Feldmann, Heike Huesmann, Stefan Eimer, Thomas Juretschke, Petra Beli, Christian Behl, Andreas Kern

## Abstract

Autophagy is a lysosomal degradation pathway that mediates protein and organelle turnover and maintains cellular homeostasis. Autophagosomes transport cargo to lysosomes and their formation is dependent on an appropriate lipid supply. Here, we show that the knockout of the RAB GTPase RAB18 interferes with lipid droplet (LD) metabolism, resulting in an impaired fatty acid mobilization. The reduced LD-derived lipid availability influences autophagy and provokes adaptive modifications of the autophagy network, which include increased ATG2B expression and ATG12-ATG5 conjugate formation as well as enhanced ATG2B and ATG9A phosphorylation. Phosphorylation of ATG9A directs this transmembrane protein to the site of autophagosome formation and this particular modification is sufficient to rescue autophagic activity under basal conditions in the absence of RAB18. However, it is incapable of enabling an increased autophagy under inductive conditions. Thus, we illustrate the role of RAB18 in connecting LDs and autophagy, further emphasize the importance of LD-derived lipids for the degradative pathway, and characterize an ATG9A phosphorylation-dependent autophagy rescue mechanism as an adaptive response that maintains autophagy under conditions of reduced LD-derived lipid availability.

## Introduction

Macroautophagy (hereafter referred to as autophagy) is a eukaryotic lysosomal degradation pathway that mediates protein and organelle turnover and maintains cellular homeostasis [1]. Autophagic cargo is transported in vesicles, so-called autophagosomes, to lysosomes for degradation and recycling of building blocks. The generation of autophagosomes is dynamic and increases within minutes upon autophagy induction. Several conditions, including nutrient deprivation or rapamycin treatment, stimulate autophagy and result in activation of the AMP-activated kinase PRKAA1 and/or inhibition of the kinase MTOR [2]. Subsequently, ULK1 kinase is activated by phosphorylation, which is a pre-requisite for the generation of autophagosomes [3].

The synthesis of double-membraned autophagosomes starts with a precursor membrane, the phagophore, which originates from specific omega-shaped domains (omegasomes) in the ER membrane [4-6]. The phagophore elongates by the addition of lipids or membranes until it finally closes to form an autophagosome, which completely incorporates the autophagic cargo. This process involves a cascade of proteins, including two ubiquitin-like conjugations systems, that mediate the formation of the ATG12-ATG5/ATG16L1 protein complex as well as the lipidation of Atg8 family members, such as MAP1LC3B (shortly LC3) [7]. Indeed, the ATG12-ATG5/ATG16L1 complex localizes to the phagophore membrane and shows E3 ligase-like activity that promotes the final step of Atg8 lipidation [8-10]. Lipid-conjugated Atg8 proteins associate with the growing phagophore membrane and are essential for autophagosome formation [11]. The proteins stay (partially) attached to autophagic vesicles, which results in their subsequent lysosomal degradation.

Besides complex protein processing, phagophore elongation is highly dependent on sufficient lipid and membrane supply [3,12]. Several cellular organelles or compartments, including ER, plasma membrane, mitochondria, Golgi complex, the ER-Golgi intermediate compartment, recycling endosomes [3,13] as well as lipid droplets [14-16], have been suggested as potent sources, donating lipids via vesicles or membrane contact sites. However, the exact process and regulation of autophagic lipid acquisition remains unresolved. A key factor is the multipass transmembrane protein ATG9A that connects the peripheral cellular compartments to autophagosome formation [17-20]. ATG9A directs vesicles from the cellular periphery to the site of phagophore maturation, where they serve as membrane sources for the buildup of autophagosomes. Additionally, these vesicles deliver essential autophagy proteins such as ATG16L1 [21,22]. Under basal autophagy conditions, the majority of ATG9A is located at the Golgi complex and switches to a dispersed vesicular localization upon autophagy induction, resulting in the enhanced shuttling of vesicles to the site of autophagosome biogenesis. Indeed, upon increased autophagy rates the specific phosphorylation of ATG9A, mediated by ULK1 and SRC kinase [23,24], is enhanced, which fosters its trafficking activity and facilitates autophagosome formation.

A further lipid source for phagophore elongation that acts independently of ATG9A, but in close functional association with the ER, are lipid droplets (LDs) [14-16]. These cytoplasmic lipid stores are found in virtually all eukaryotic cells and are composed of an organic phase of neutral lipids, mainly triacylglycerides (TAGs) and sterol esters, which are separated from the cytosol by a phospholipid monolayer and coated by a distinct network of proteins [25]. Under lypolytic conditions, TAGs are hydrolyzed by the activity of various lipases and the generated fatty acids are employed for energy metabolism or modified to appropriate lipids within the ER [26]. Importantly, recent studies, employing yeast or human cell lines, have demonstrated that specific LD-associated lipases, including PNPLA2/ATGL and PNPLA5, modulate autophagy and that LD-derived lipids support the efficient formation of autophagosomes [14-16,27].

The RAB GTPase RAB18 has functionally been associated with LD metabolism [28] and is a positive modulator of autophagy [29]. RAB18 localizes to LDs in adipocytes, fibroblasts, epithelial cells, and several clonal cell lines. This association has been connected to a function in LD homeostasis and the release of stored fatty acids [28,30,31]. The interaction of RAB18 with ER-linked tethering factors mediates the connection of LDs with the ER membrane and facilitates LD homeostasis [32,33]. In a human mammary carcinoma cell line, however, a RAB18 function in LD biogenesis or turnover has not been observed [34]. Importantly, loss-of-function mutations in *RAB18* cause Warburg Micro syndrome (WARBM) [35,36], a severe human autosomal recessive (neuro-)developmental disorder. The molecular mechanisms responsible for the pathology remain unclear, yet, WARBM patient cells are characterized by an altered appearance of LDs [37].

In this study, we examined the impact of the stable RAB18 knockout on LD metabolism and autophagy in HeLa cells. The permanent loss of the RAB GTPase provoked a LD phenotype that strongly resembled WARBM patient cells. In depth analysis revealed, that LDs were enlarged in size and reduced in number and the release of fatty acids was impaired in the absence of RAB18. We observed that insufficient LD-derived lipid availability influenced autophagy, resulting in adaptive adjustments of the autophagy network, which included an increased expression and altered phosphorylation of ATG2B and the enhanced generation of ATG12-ATG5 conjugates. Most strikingly, the phosphorylation of ATG9A at amino acid residues Y8 and S14, which direct the protein to the autophagic pathway, were enhanced in RAB18 KO cells. This modification rescued autophagy under basal autophagy conditions. Thus, this study strengthens the function of RAB18 in LD metabolism and further demonstrates the importance of LD-derived lipids for autophagy. Moreover, we characterize an ATG9A phosphorylation-dependent rescue mechanism as a potent adaptive response to maintain basal autophagic activity under conditions of deficient LD-derived lipid availability.

## Results and Discussion

In previous studies, using transient genetic manipulations, we have characterized RAB18 as a positive modulator of autophagy [29]; reduced protein levels of the RAB GTPase decreased autophagic activity while increased levels enhanced the degradative pathway. To conduct a more detailed analysis of RAB18 function in autophagy, we generated stable RAB18 KO HeLa cells, employing the CRISPR/CAS9 technology, and selected two independent clonal cell lines (Fig. S1).

Initially, we focused on LDs since RAB18 has functionally been linked to their metabolism and an altered LD phenotype has been described for WARBM patient cells [30,37]. Fluorescence imaging of LDs revealed a striking LD appearance in the absence of RAB18 (Fig. 1A), similar to WARBM patient cells [37]. LDs accumulated in the perinuclear region and their detailed analyses demonstrated that they were enlarged in size and reduced in number per cell, when compared to HeLa wild type (WT) cells (Fig. 1B+C). This finding was confirmed by transmission electron microscopy showing fewer but enlarged LDs, which are mostly localized close to the nucleus (Fig. 1D).

**Figure 1.**
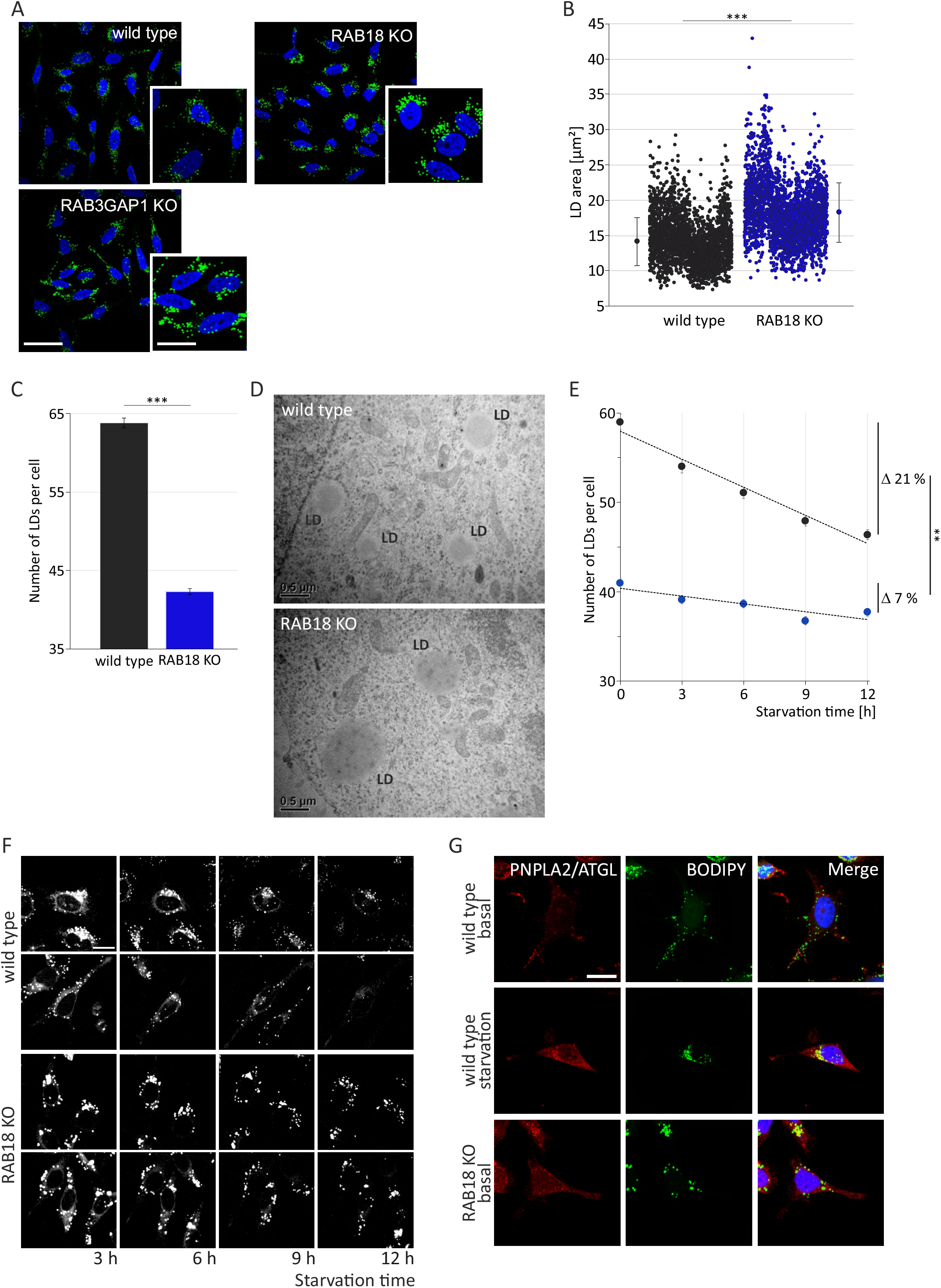
The loss of RAB18 interferes with LD homeostasis and mobilization of fatty acids. **A** Representative confocal fluorescence images of LDs in wild type, RAB18 KO and RAB3GAP1 KO cells stained with BODIPY 493/505. Scale bar: 50 and 100 µm. **B** Analyses of LD area in approx. 2500 single wild type and RAB18 K.O cells. Statistics are depicted as mean ± SD. n = 3, *t*-test, ***P ≤ 0.001. **C** Analyses of LD number in approx. 2500 single wild type and RAB18 KO cells. Statistics are depicted as mean ± SEM. n = 3, *t*-test, ***P ≤ 0.001. **D** Transmission electron microscopic image of LDs in wild type and RAB18 KO cells. **E** LD consumption upon starvation in wild type (grey) and RAB18 KO cells (blue). At each indicated time point during the starvation period, the number of LDs per cell was evaluated in approx. 1350 wild type and RAB18 KO cells. Statistics are depicted as mean ± SEM. n = 3, *t*-test, **P ≤ 0.01. **F** Confocal fluorescence life cell images of wild type and RAB18 KO cells pre-treated with BODIPY 558/568 C_12_ and imaged during EBSS treatment. The depicted cells are representative for 3 independent approaches. Scale bar: 20 µm. **G** Representative confocal fluorescence images of wild type and RAB18 KO cells stained with PNPLA2/ATGL (red) and BODIPY 493/505 (green). Cell nuclei are stained with DAPI (blue). Wild type cells are shown under basal and starvation conditions (2 h EBSS treatment). Scale bar: 20 µm.

Importantly, we observed an identical LD phenotype in CRISPR/CAS9-mediated stable RAB3GAP1 KO cells (Fig. 1A). RAB3GAP1 acts as an upstream regulator of RAB18 activity [29,33,38] and loss-of-function mutations in *RAB3GAP1* have likewise been associated with WARBM [39]. Thus, we detected an altered LD appearance in cell lines with a KO in different - but functionally associated - genes, which emphasizes the relevance of RAB18 and RAB3GAP1 for LD homeostasis.

Presuming that the LD phenotype in KO cells originates from an altered LD metabolism, we investigated the consumption of the lipid stores under nutrient-deprived conditions. Nutrient deprivation results in the rapid hydrolysis of TAGs and the use of released fatty acids for mitochondrial energy metabolism [40]. Indeed, starvation triggered a time-dependent reduction in LD numbers in WT cells, while this process was repressed in RAB18 KO cells (Fig. 1E). The number of LDs per cell showed only a marginal decline during the starvation period. Hence, the absence of RAB18 interfered with the turnover of LDs.

In order to directly investigate the mobilization of fatty acids from LDs upon starvation, we monitored their release, employing the dye-conjugated fatty acid BODIPY 558/568 C_12_ and life cell imaging (Fig. 1F). As expected, in WT cells, BODIPY 558/568 C_12_ efficiently migrated out of LDs under nutrient-deprived conditions; the LD-located fluorescence signal decreased and the area of single LDs progressively shrunk during the starvation period. In contrast, the dye-conjugated fatty acid was mainly immobile in RAB18 KO cells and remained within LDs during the starvation period, confirming that fatty acid mobilization in RAB18 KO cells is indeed disturbed and is responsible for the LD phenotype.

We further examined the role of RAB18 in the formation of LDs and monitored their numbers after the supplementation with oleic acid. Fatty acid treatment results in the rapid buildup of LDs and, noteworthy, we observed no differences comparing RAB18 KO and WT cells (Fig. S2). This indicates that the induced generation of LDs was not affected by the stable loss of RAB18.

Another key component of LD metabolism is the lipase PNPLA2/ATGL, which is responsible for the hydrolysis of TAGs [41,42]. Under lypolytic conditions, the enzyme localizes to LDs via its membrane-embedded target sequence and mediates the rapid release of fatty acids [41]. Excitingly, the TAG lipase has also been characterized as a modulator of autophagy [27]. We analyzed the localization of PNPLA2/ATGL at LDs in RAB18 KO and WT cells, employing immunocytochemistry (Fig. 1G). Interestingly, the lipase accumulated at LDs in RAB18 KO cells, demonstrating that the loss of RAB18 did not modulate its transfer to the surface of the lipid stores. However, the enhanced LD-localization was not associated with an efficient fatty acid release, emphasizing that LD turnover is impaired in the absence of RAB18.

In summary, so far we found that the loss of RAB18 function caused a defect in the consumption of LDs, producing the prominent LD phenotype; a clear indication that cellular lipid metabolism and, in particular, LD-derived lipid availability is disturbed in the absence of the RAB GTPase.

Next, we analyzed the influence of the observed alterations in LD metabolism on autophagy, considering that the degradative pathway is strongly dependent on an adequate lipid supply. Intriguingly, immunoblot analyses of autophagic activity in stable RAB18 KO cells showed that the loss of RAB18 did not affect basal autophagy (Fig. 2A). Flux analyses of the lipidated form of LC3, LC3II, showed no differences comparing the cell lines under basal, uninduced, autophagy conditions. This result was supported by fluorescence microscopic investigations that examined the total number of LC3- and SQSTM1-positive autophagosomes (Fig. 2B). The amount of autophagosomal structures was comparable in RAB18 KO and WT cells under basal conditions.

**Figure 2.**
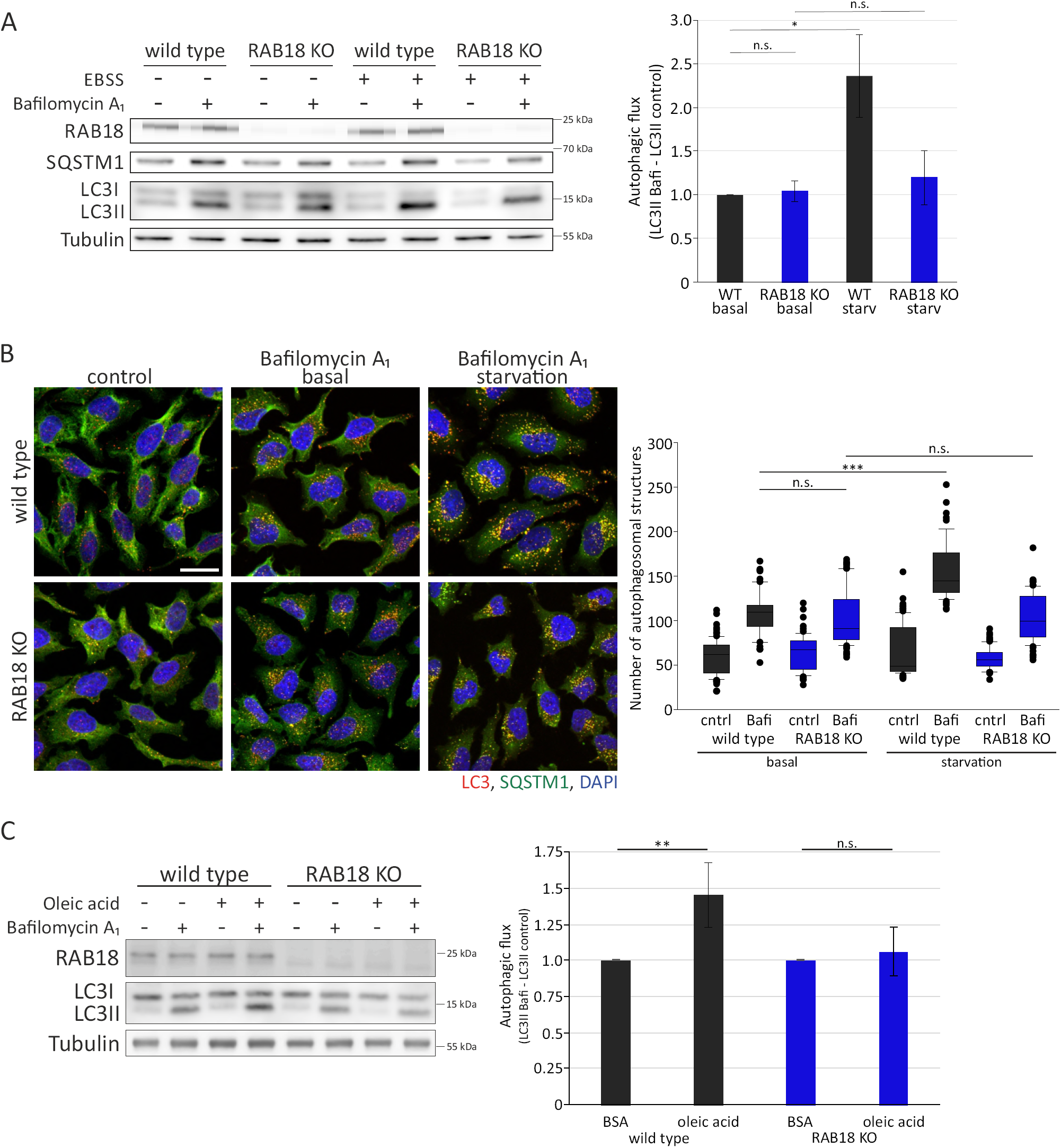
The effect of reduced LD-derived lipid availability on autophagy in RAB18 KO cells. **A** Immunoblot analyses of autophagic activity in wild type and RAB18 KO cells under basal and induced autophagy conditions (2 h EBSS treatment). Cells were treated with DMSO (control) or Bafilomycin A_1_ for 2 h to allow LC3II flux evaluation. Statistics are depicted as mean ± SD. n = 4, *t*-test, *P ≤ 0.05. **B** Immunocytochemical stainings of LC3 (red) and SQSTM1 (green) in wild type and RAB18 KO cells. Cell nuclei are stained with DAPI (blue). Cells were treated with DMSO (control) or Bafilomycin A_1_ under control conditions or EBSS treatment. For statistics autophagosomal structures were quantified in 60 - 80 single cells per cell line and treatment from 3 independent experiments, *t*-test, ***P ≤ 0.001. **C** Immunoblot analyses of autophagic activity after oleic acid pre-treatment. Wild type and RAB18 KO cells were pre-treated with BSA (control) or BSA-conjugated oleic acid (14 h) and thereafter with DMSO (control) or Bafilomycin A_1_ for 3 h to allow LC3II flux evaluation. Statistics are depicted as mean ± SD, *t*-test, **P ≤ 0.01.

Subsequent starvation experiments induced autophagy in WT cells, however, excitingly, had no effect on autophagic activity in RAB18 KO cells: the cells did not respond to nutrient deprivation with the usually observed increased autophagic flux (Fig. 2A+B). Importantly, we confirmed this particular finding in RAB3GAP1 KO cells (Fig. S3); the stable KO of RAB3GAP1 had no impact on basal autophagy, but impeded the induction-mediated increased autophagic activity.

These results obtained in stable HeLa knockout cells actually opposed our findings gained by transient knockdown of RAB18 as well as RAB3GAP1 [29,43]; siRNA-mediated reduction of protein levels caused a decline in autophagic activity, both, under basal as well as induced autophagy conditions. The novel findings presented here indicate that the stable RAB18 (as well as RAB3GAP1) KO lead to an adaptation of the autophagy network, which maintains basal autophagic activity and compensates the permanent loss of RAB18 function. However, this adaptive response is incapable of allowing enhanced autophagy upon induction.

The prominent defect in RAB18 KO cells is the reduced fatty acid transfer from LDs, which results in an insufficient lipid supply to support autophagosome formation. Several publications have emphasized the link between LD-derived lipids and autophagy [14-16]. A recent study has reported that the increased accessibility of fatty acids from LDs enhances autophagic capacity by facilitating autophagosome formation and, importantly, that this process is dependent on LD-located lipases [14]. To analyze the significance of LD-derived lipids for autophagy in our system, we investigated autophagic activity after pre-treatment with oleic acid. The resulting enhanced availability of LD-derived lipids stimulated autophagy in WT cells and facilitated the flux of LC3II (Fig. 2C), confirming the previous study [14]. Strikingly, these conditions did not affect autophagic capacity in RAB18 KO cells. The pre-treatment with oleic acid did not result in an increased autophagic activity. This data strongly indicates that the transfer of fatty acids from LDs to the autophagic pathway is disturbed in the absence of RAB18 and further emphasizes the importance of LD-derived lipids for autophagy.

Considering that basal autophagy was not affected by the loss of RAB18, the results obtained so far pointed towards an adaptation of the autophagy network to conditions of reduced LD-derived lipid availability. In order to analyze potential adaptive adjustments, we investigated the status of the canonical autophagy induction pathway and analyzed phosphorylation of MTOR and ULK1 (Fig. 3). MTOR phosphorylation at amino acid residue S2448 regulates the kinase [44]. Indeed, we observed that S2448 phosphorylation levels were decreased under autophagy-induced conditions and, interestingly, we found no difference in total levels of this specific MTOR modification comparing RAB18 KO and WT cells (Fig. 3A, Fig. S4). The kinase ULK1 is phosphorylated at multiple sites to regulate its activity, therefore, we examined the MTOR-mediated, inactivating, phosphorylation at S758 as well as the PRKAA1-mediated, activating, phosphorylation at S555 [45,46]. Interestingly, total levels of both post-translational modifications were comparable in both cell lines, which emphasizes that ULK1 activity is not altered by the loss of RAB18 (Fig. 3A, Fig. S4). Additionally, we conducted a SILAC-based quantitative phosphoproteomics approach that did not identify alterations in MTOR and ULK1 phosphorylation at these residues (data not shown). These findings indicate, that possible adjustments of the autophagy network, which functionally compensate the loss of RAB18, are not related to alterations in the activity of the canonical autophagy induction system.

**Figure 3.**
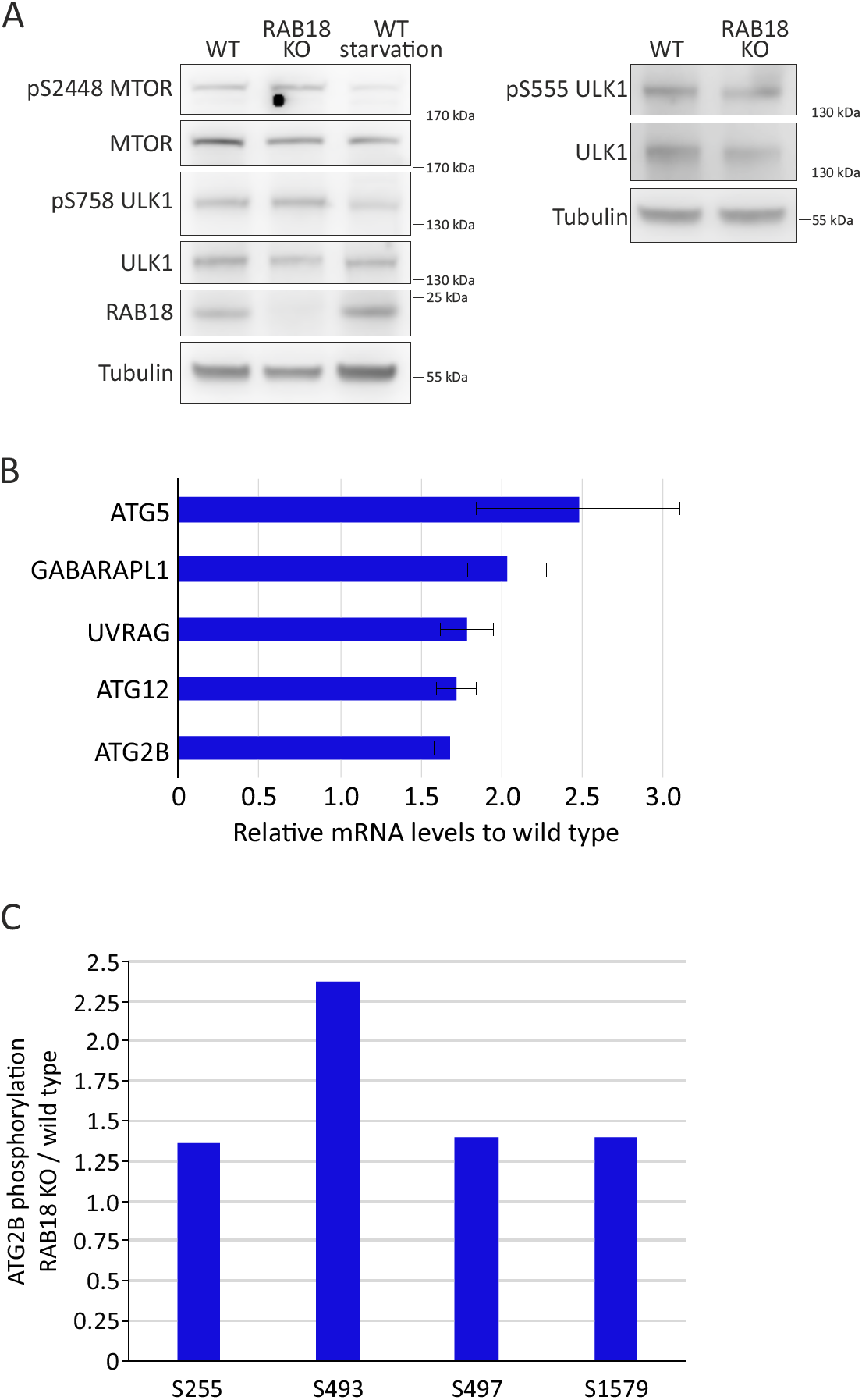
Analysis of the autophagy network in RAB18 KO cells. **A** Immunoblot analyses of MTOR and ULK1 phosphorylation in wild type and RAB18 KO cells under basal and induced conditions (2 h EBSS treatment). **B** Representation of genes showing increased expression levels in RAB18 KO compared to wild type cells as evaluated using the qPCR array in 3 independent approaches. The threshold for upregulation was set to a 1.5-fold increase mRNA levels. **C** Phosphorylation levels of ATG2B (Q96BY7) serine residues in RAB18 KO compared to wild type cells, determined by SILAC-based mass spectrometry and depicted as mean values of 2 replicates.

Moreover, we investigated expression levels of key autophagy proteins, employing a qPCR array and found that mRNA levels of a discrete number of autophagy factors were up-regulated in RAB18 KO cells (Fig. 3B; Table S1). Remarkably, all these proteins can be allocated to early steps of the autophagy pathway, suggesting that the network responded to the stable protein loss with adaptations that facilitate autophagosome formation. Here, the increased expression of ATG2B is of particular interest. Mammalian cells possess two functionally redundant ATG2 proteins, ATG2A and ATG2B, which are suggested to regulate phagophore growth and support the closure of autophagosomes [47,48]. Noteworthy, ATG2A/B have been described to localize to phagophores as well as LDs and their knockdown also results in the accumulation of enlarged LDs [47,49], indicating that ATG2 proteins might be involved in linking LD metabolism and autophagy. Thus, we presume that the up-regulation of ATG2B is a direct adaptation to a disturbed LD-derived lipid transfer. Furthermore, the quantitative phosphoproteomics approach detected several phosphorylation sites of ATG2B and revealed that solely phosphorylation of the amino acid residue S493 was considerably enhanced in the absence of RAB18 (Fig. 3C). This phosphorylation site has not been functionally characterized yet, but the increased ATG2B expression in conjunction with the altered phosphorylation suggests that ATG2B function is relevant for the adaptive response of the autophagy network.

Within the autophagy system, ATG9A is well-acknowledged to function in the transfer of vesicles from the cellular periphery to the site of autophagosome formation to supply required membranes [17,20,22]. Notably, mRNA and protein levels of ATG9A were not altered in RAB18 KO compared to WT cells (Fig. 4B, Fig. S5). However, it has recently been shown that the autophagic activity of ATG9A is actually regulated by phosphorylation at amino acid residues S14 and Y8 [23,24]. In consequence, we analyzed the ULK1-mediated phosphorylation of ATG9A at S14, employing the quantitative phosphoproteomics approach. Indeed, the level of this particular phosphorylation of ATG9A was slightly but reproducibly enhanced in RAB18 KO cells (Fig. 4A). Thus, even though we did not find any clear indication of an altered ULK1 activity (Fig. 3A), the ULK1-mediated phosphorylation of ATG9A was enhanced in the absence of RAB18.

**Figure 4.**
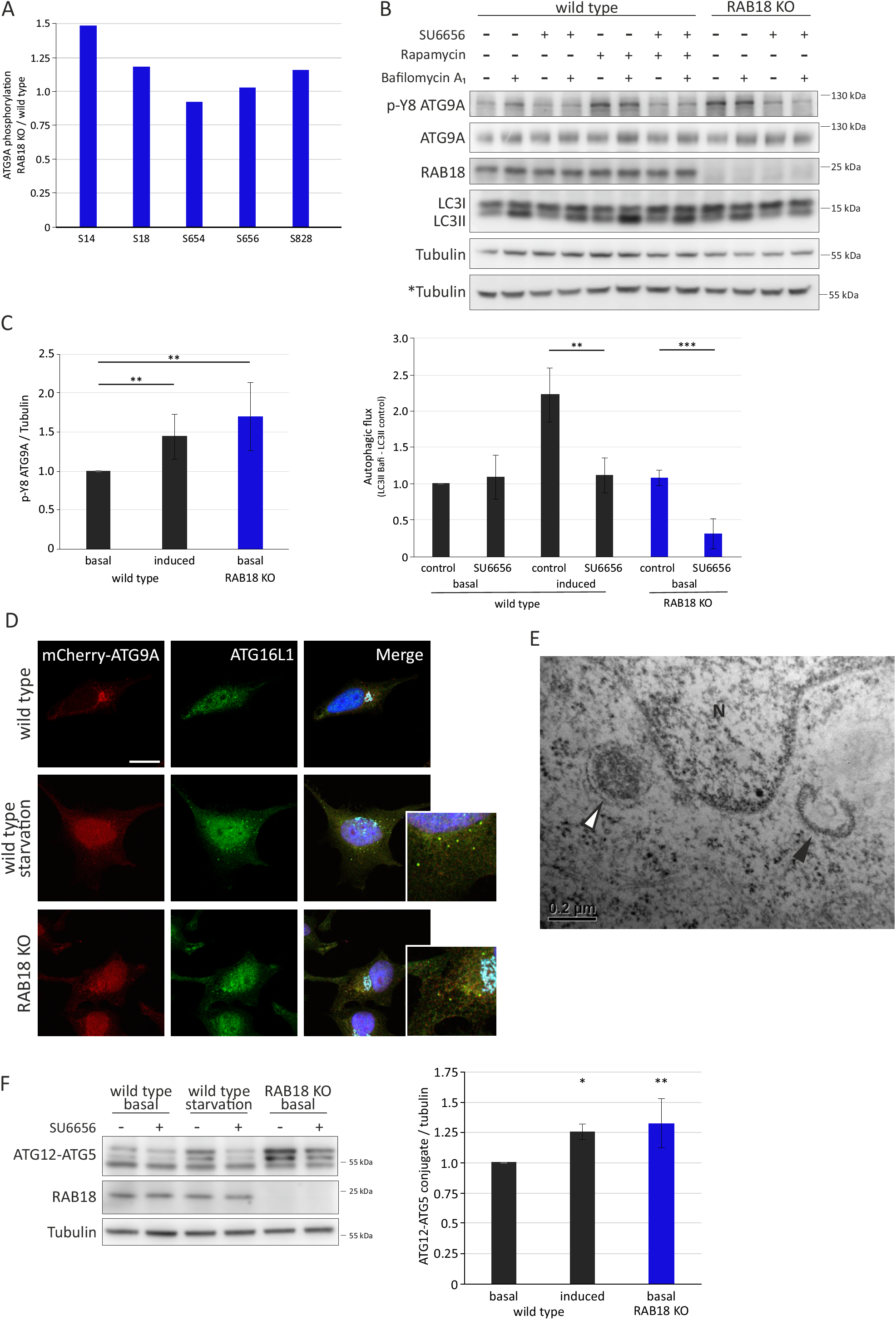
RAB18 KO-mediated adaptations of the autophagy network. **A** Phosphorylation levels of ATG9A (Q7Z3C6) serine residues in RAB18 KO compared to wild type cells, determined by SILAC-based mass spectrometry and depicted as mean values of 2 replicates. **B** Immunoblot analyses of p-Y8 ATG9A levels and autophagic activity in wild type and RAB18 KO cells. Inhibition of SRC kinase was mediated by pre-treatment with SU6656 for 12 h. Wild type cells were maintained in full medium (basal) or transferred into EBSS (2h) for autophagy induction. Cells were treated with DMSO (control) or Bafilomycin A_1_ for 2 h to allow LC3II flux evaluation. *indicates Tubulin loading control only corresponding to parallel ATG9A blot. **C** Statistical analyses of B depicted as mean ± SD. n = 5, *t*-test, **P ≤ 0.01, ***P ≤ 0.001. **D** Representative confocal fluorescence images of mCherry-ATG9A (red), ATG16L1 (green), GM130 (Golgi, turquois), and DAPI (nuclei, blue) of wild type and RAB18 KO cells. Wild type cells were treated for 3 h with EBSS (starvation). Scale bar: 20 µm. **E** Transmission electron microscopic image of pre-autophagosomal structures in RAB18 KO cells under basal autophagy conditions. Open arrowhead: omegasome; filled arrowhead: phagophore; N: nucleus. **F** Immunoblot analyses of ATG12-ATG5 conjugate levels in wild type and RAB18 KO cells. Inhibition of SRC kinase was mediated by pre-treatment with SU6656 for 12 h. Wild type cells were maintained in full medium (basal) or transferred into EBSS (2h) for autophagy induction. Statistics are depicted as mean ± SD. n = 4, *t*-test, *P ≤ 0.05, **P ≤ 0.01.

Since the quantitative phosphoproteomics approach solely detected the serine residues on ATG9A, we investigated phosphorylation of the Y8 residue of ATG9A (referred to as p-Y8 ATG9A) by immunoblotting. Consistent with a recent study [24], we found that p-Y8 ATG9A levels were increased upon autophagy induction in WT cells (Fig. 4B+C). This correlated with the expected enhanced autophagic activity. The use of a specific SRC kinase inhibitor (SU6656) efficiently prevented Y8-phosphorylation and impeded the enhanced autophagic activity. This confirms the functional link between this particular SRC kinase-dependent phosphorylation of ATG9A and autophagic activity. Most strikingly, p-Y8 ATG9A levels were already enhanced under basal conditions in RAB18 KO cells, which, importantly, did not correspond to an increased autophagic activity in this case (Fig. 4 B+C). Remarkably, the inhibition of Y8-phosphorylation significantly decreased basal autophagic activity in RAB18 KO cells. This strongly argues that increased p-Y8 ATG9A levels are indeed causative for the maintenance of basal autophagy in the absence of RAB18.

ATG9A is prominently located at the Golgi complex and changes its cellular localization to a dispersed vesicular distribution upon autophagy induction [18,20,21]. This leads to enhanced vesicle trafficking to the site of autophagosome formation and is directly linked to increased numbers of perinuclear ATG16L1-positive pre-autophagosomal structures [21,22]. Employing confocal fluorescence microscopy, we analyzed the localization of ATG9A in RAB18 KO and WT cells under basal and induced autophagy conditions (Fig. 4D). As expected, the majority of ATG9A was located at the Golgi complex in WT cells, but changed its localization to a dispersed vesicular distribution upon starvation-mediated autophagy induction. This altered localization of ATG9A was indeed accompanied by increased numbers of ATG16L1 punctae, which were mostly ATG9A positive. Importantly, RAB18 KO cells showed the same dispersed vesicular distribution of ATG9A, which was also associated with enhanced numbers of ATG16L1-positive structures, already under basal autophagy conditions. In accordance with this observation, transmission electron microscopy revealed an increased number of perinuclear pre-autophagosomal structures, such as omegasomes and phagophores (Fig. 4E), which are not seen in WT cells.

Summing up, the analyses of ATG9A phosphorylation and localization clearly illustrate that the autophagic activity of ATG9A is enhanced in the absence of RAB18, which is an essential alteration to maintain basal autophagy under conditions of deficient LD-derived lipid availability.

Since RAB18 KO cells are characterized by basal autophagy network adaptations that resemble induced autophagy conditions and, in addition, we found that expression levels of ATG12 and ATG5 were increased (Fig. 3B), we consequently also analyzed the ATG12-ATG5 conjugate formation. Both proteins are covalently linked by an ubiquitin-like conjugation process [10,50], which is a pre-requisite for the formation of the ATG12-ATG5/ATG16L1 complex that is essential for the lipidation of Atg8 family members [10]. Employing immunoblotting, we observed that total ATG12-ATG5 conjugate levels were increased in RAB18 KO cells under basal autophagy conditions and resembled ATG12-ATG5 levels of WT cells after induction (Fig. 4F). Thus, the increased trafficking of ATG9A to the site of autophagosome formation is associated with an enhanced (autophagy induction-like) generation of ATG12-ATG5 conjugates in RAB18 KO cells already under basal autophagy conditions. We additionally observed that the formation of the ATG12-ATG5 conjugate is dependent on SRC kinase activity, since the treatment with the SRC kinase inhibitor potently reduced total conjugate levels (Fig. 4F). It is tempting to speculate, that ATG12-ATG5 conjugate levels are indeed directly dependent on ATG9A activity.

The loss of RAB18 and the associated insufficient lipid supply for autophagosome formation result in adaptive adjustments of the autophagy network, which resemble alterations that are typically observed in WT cells upon autophagy induction. These adaptations are sufficient to maintain basal autophagy, but incapable of enabling a further increase in autophagic activity upon nutrient deprivation. We conclude that the SRC kinase-dependent phosphorylation of ATG9A at Y8 is the critical rescue mechanism to maintain basal autophagy under conditions of deficient LD-derived lipid availability (Fig. 5). In summary, this emphasizes the importance of LD-derived lipids for autophagy and further illustrates a detailed molecular function of RAB18, which connects LD metabolism and autophagy. Furthermore, these findings may reflect on a potential pathogenetic process for the development of WARBM.

**Figure 5.**
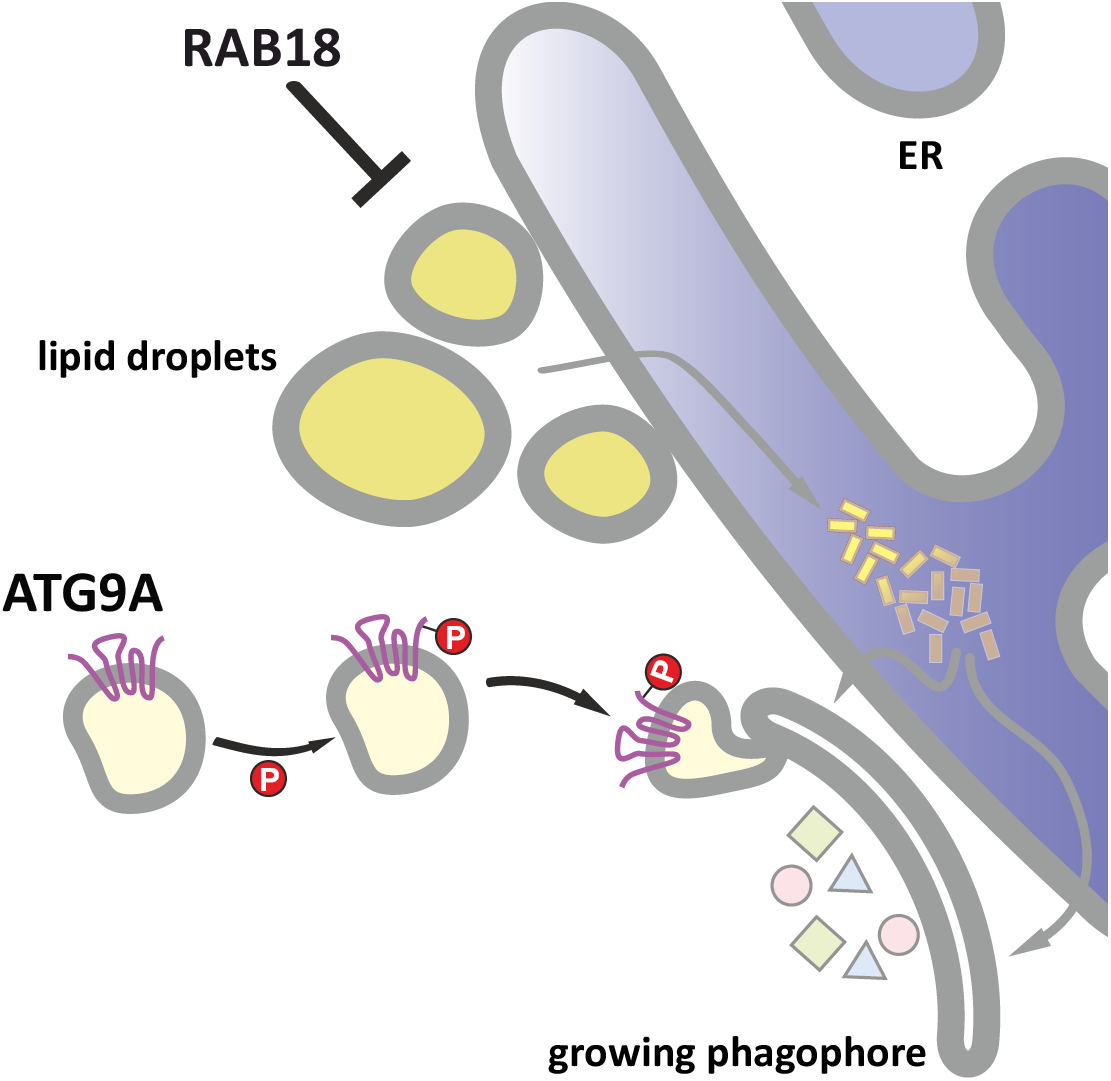
Schematic synopsis. The loss of RAB18 function impedes the mobilization of fatty acids form LDs to the ER, which results in an insufficient transfer of lipids to the site of autophagosome formation. This provokes an adaptive response of the autophagy network, including the enhanced phosphorylation of ATG9A. The resulting augmented ATG9A-mediated vesicle transport rescues basal autophagy, but is incapable of enabling a further increase in autophagic activity under induced conditions.

## Material and Methods

### Cell culture

HeLa cells were cultured in DMEM, supplemented with 10 % fetal bovine serum, 1 mM sodium pyruvate, and 1 x antibiotic-antimycotic solution at 37 °C in a 5 % CO_2_-humidified atmosphere. Stock solutions of Bafilomycin A_1_ (Biozol, TRC-B110000) and Rapamycin (Enzo, BML-A275-0025) were prepared in DMSO and employed as described before [43]. Stock solutions of SU6656 SRC Inhibitor (Selleck Chemicals, S7774) were prepared in DMSO and cells were treated with 10 µM for 12 h. Expression plasmid encoding mCherry-ATG9A was kindly provided by Ivan Dikic (Frankfurt, Germany). Cells were transfected with 10 µg of plasmid. All transfections were carried out by electroporation as described previously [43].

### CRISPR/CAS9 nuclease RNA-guide target deletions

Gene specific CRISPR/CAS9 vectors targeting *RAB18* and *RAB3GAP1* were obtained from Sigma Aldrich. Vectors: pLV-U6g-EPCG-RAB18-I gRNA: AATAGTCCAATCCTGAAGTGGG; pLV-U6g-EPCG-RAB18-II gRNA: GTGCACCTCTATAATAGCTGGG; pLV-U6g-EPCG-RAB3GAP1-I gRNA: TCCGAGGTATTTGAGATCACGG; pLV-U6g-EPCG-RAB3GAP-II gRNA: TATGGGCTACGTGAGTTCG TGG. Wild type HeLa cells were transfected with 15 μg of plasmids and selection of positive clones was performed using culture medium containing 1 μg/ml puromycin. Positive knockout clones were identified via immunoblotting and qPCR.

### Immunoblotting

Immunoblot analyses were performed as previously described [29]. Usually 15 μg of total protein were subjected to hand-cast 12 % Bis-Tris or 4-12 % NuPage Novex Bis-Tris gels (Thermo Scientific, MP0335) and transferred onto nitrocellulose membranes. Membranes were blocked with 5 % (w/v) milk powder in TBS containing 0.05 % Tween 20 for 1.5 h at RT and afterwards probed with the appropriate primary (1:500; 4 °C o.N.) and secondary antibodies (1,5 h at RT). Proteins were detected by chemiluminescence and developed using the Amersham Imager 600 (GE Healthcare Life Science). Primary antibodies used in this study: MAP1LC3B (Sigma, L7543), SQSTM1 (Progen, GP62-C), RAB18 (Proteintech, 11304-1-AP), RAB18 (Santa Cruz Biotechnology, sc-393168), RAB3GAP1 (Sigma, SAB4500914), RAB3GAP2 (Sigma, HPA026273), Tubulin (Sigma, T9026), ATGL/PNPLA2 (Cell Signaling, 2138), ATG5 (Novus Biologicals, NB110-53818), ATG9A (Cell Signaling, 135095), p-Y8 AT9A (kindly provided by Yushan Zhu and Quan Chen, Nankai, China), ULK1 (Cell Signaling, 8054), p-ULK1 Ser555 (Cell Signaling, 5869), p-ULK1 Ser757 (Cell Signaling, 6888), mTOR (Cell Signaling, 2972S), p-mTOR S2448 (Cell Signaling, 2971).

### Immunocytochemistry

Cells were grown on glass cover slips, washed with PBS and fixed with 4 % (w/v) PFA for 20 min at RT. Cells were permeabilized in 90 % ice-cold methanol for 6 minutes or 0.02 % Triton X-100 for 20 minutes at RT. Unspecific binding sites were blocked with 3 % (w/v) BSA in PBS for 1 h at RT. Subsequently, cells were incubated with primary antibodies (1:200 in 1 % (w/v) BSA in PBS) for 1 h at RT, and thereafter with fluorophore-conjugated secondary antibodies (1:200 in PBS) followed by DAPI. Primary antibodies used: ATGL/PNPLA2 (Cell Signaling, 2138), ATG16L1 (MBL, PM040), GM130 (BD biosciences, 610823), MAP1LC3B (nano Tools, 0260-100), SQSTM1 (Progen, GP62-C), Tubulin (Sigma-Aldrich, T9026). Stock solution of BODIPY 493/503 (Thermo Scientific, D3922) was prepared in ethanol according to the supplier and LDs were stained at a dilution of 1:1000. BODIPY 558/568 C_12_ (Thermo Scientific, D3835) stock solution was prepared according to the supplier; cells were probed at 5 nM for 24 h and incubated in fresh medium for 1 h before microscopy. Cells were imaged using the laser scanning microscope LSM710 (Zeiss). Imaging for quantification of autophagosomal structures was performed via the Opera Phenix High-Content Screening System (Perkin-Elmer) and the number of LC3- and SQSTM1-positive structures was quantified using Harmony High Content Imaging and Analysis software (Perkin-Elmer).

### Lipid droplet analysis

For LD analyses, control and EBSS treated cells were stained with Tubulin, DAPI and BODIPY 493/503 as described above. Cells were imaged via the Opera Phenix High-Content Screening System (Perkin-Elmer). 2650 or 1350 single cells per cell line or treatment were used for statistical analyses. Single cells and cell volume were defined via DAPI and tubulin staining, respectively. Thereafter, LD number and size were quantified by BODIPY 493/503 fluorescence, employing Harmony High Content Imaging and Analysis software (Perkin-Elmer).

### Autophagic activity analysis

Autophagic activity was analyzed in accordance with [51]. Cells were treated with Bafilomycin A_1_ for 2 - 4 h or DMSO for control and the fluxes of autophagic substrates were calculated by the subtraction of accumulated levels upon Bafilomycin A_1_ treatment and DMSO control.

### Quantitative real-time PCR

RNA extraction, reverse transcription and quantitative real time PCR were performed as described previously [43]. Primer sequences for the autophagy array are listed in Supplemental Table S1.

### Electron microscopy

For transmission electron microscopy (TEM) analysis cells were grown on 7.8 mm thick Aclar discs (Ted Pella Inc.) and fixed by immersion in 2.5 % glutaraldehyde in 0.1 M sodium cacodylate buffer pH 7.4 for 1 h at RT. After the fixation, discs were washed in 0.1 M cacodylate buffer. After treatment with 1 % OsO_4_, the sections were stained with uranyl acetate, dehydrated and flat-embedded in epon resin (Araldite 502). 35 nm ultrathin cross sections were cut using an Ultracut S ultramicrotome (Reichert) and analyzed using a CM12 TEM (Philips) operated at 80 kV and equipped with an ES500W Erlangshen (782) CCD camera (Gatan).

### Quantitative phosphoproteome analysis

Quantitative phosphoproteome analysis was performed as described previously [52]. Briefly, cells were cultured in SILAC media containing either L-arginine and L-lysine, L-arginine [^13^C_6_] and L-Lysine [^2^H_4_], or L-arginine [^13^C_6_-^15^N_4_] and L-lysine [^13^C_6_-^15^N_2_] (Cambridge Isotope Laboratories). Cells were lysed in modified RIPA buffer (50 mM Tris pH 7.5, 650 mM NaCl, 1 mM EDTA, 1 % NP-40, 0.1 % sodium deoxycholate) supplemented with protease and phosphatase inhibitors. Proteins were precipitated in ice-cold acetone and re-dissolved in denaturation buffer (6 M urea, 2 M thiourea in 10 mM HEPES pH 8.0). Cysteines were reduced with 1 mM dithiothreitol and alkylated with 5.5 mM chloroacetamide. Proteins were digested with endoproteinase Lys-C (Wako Chemicals) and sequencing grade-modified trypsin and thereafter purified using reversed-phase Sep-Pak C18 cartridges (Waters). Phosphorylated peptides were enriched and fractionated using micro-column-based strong-cation exchange chromatography and desalted on reversed-phase C18 StageTips. Peptide fractions were analyzed on a quadrupole Orbitrap mass spectrometer (QExactive Plus, Thermo Scientific) equipped with a UHPLC system (EASY-nLC 1000, Thermo Scientific). Survey full-scan MS spectra were acquired in the Orbitrap. The ten most intense ions were sequentially isolated and fragmented by higher energy C-trap dissociation (HCD). Fragment spectra were acquired in the Orbitrap mass analyzer. Raw data files were analyzed using MaxQuant (dev. version 1.5.2.8). Parent ion and MS2 spectra were searched against a database containing 95,057 human protein sequences obtained from the UniProtKB released in May 2018 using Andromeda search engine. Site localization probabilities were determined by MaxQuant using the PTM scoring algorithm as described previously [53]. The dataset was filtered based on posterior error probability to arrive at a false discovery rate below 1 % estimated using a target-decoy approach [54]. Only phosphorylated peptides with a score ≥40, delta score ≥8, score difference ≥5 and localization probability ≥0.75 were considered for downstream analysis.

### Statistics

Statistical significance was determined by Student’s *t*-test or Mann-Whitney *U*-test in dependence of the normal distribution or variance differences of the samples using SIGMA STAT (SPSS Science). Statistical significance was accepted at a level of P < 0.05. The results are expressed as mean ± standard deviation (SD) or standard error of the mean (SEM).

## Acknowledgements

We would like to thank Yushan Zhu and Quan Chen (Nankai, China) for kindly supplying the ATG9A phospho-specific antibody and Sandra Ritz (IMB, Mainz, Germany) for her support with the life cell imaging. We also thank Marion Basoglu from the Electron Microscopy Facility of the Biology department for her help with the TEM sample preparation and imaging. This work is supported by the Collaborative Research Center CRC1177 of the DFG (to CB and PB), the Heller foundation (to CB) and the Emmy Noether Program BE 5342/1-1 of the DFG (to PB).

## Author contributions

FB, DS, AF, HH carried out the experiments. SE conducted the electron microscopy. TJ and PB performed the phosphoproteomics analysis. FB, DS, PB, CB and AK analyzed data. CB and AK designed the study and wrote the manuscript with contribution from FB and DS.

---

**Figure S1.**
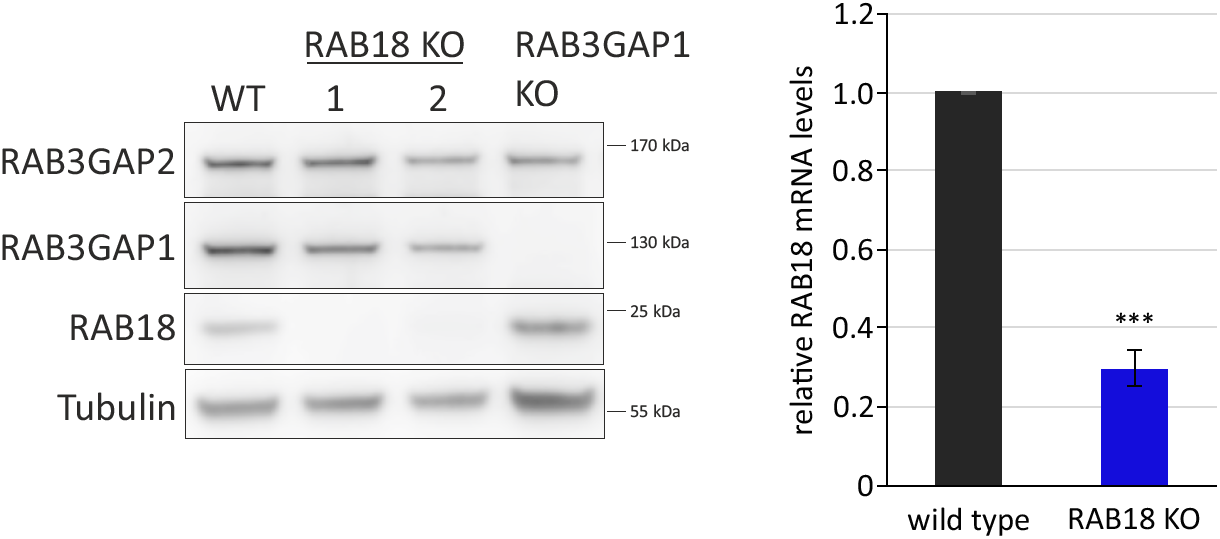
Characterization of knockout. Immunoblot and qPCR analyses of RAB18 protein and mRNA levels as well as RAB3GAP1 protein levels in the selected knockout clones in comparison to HeLa wild type cells.

**Figure S2.**
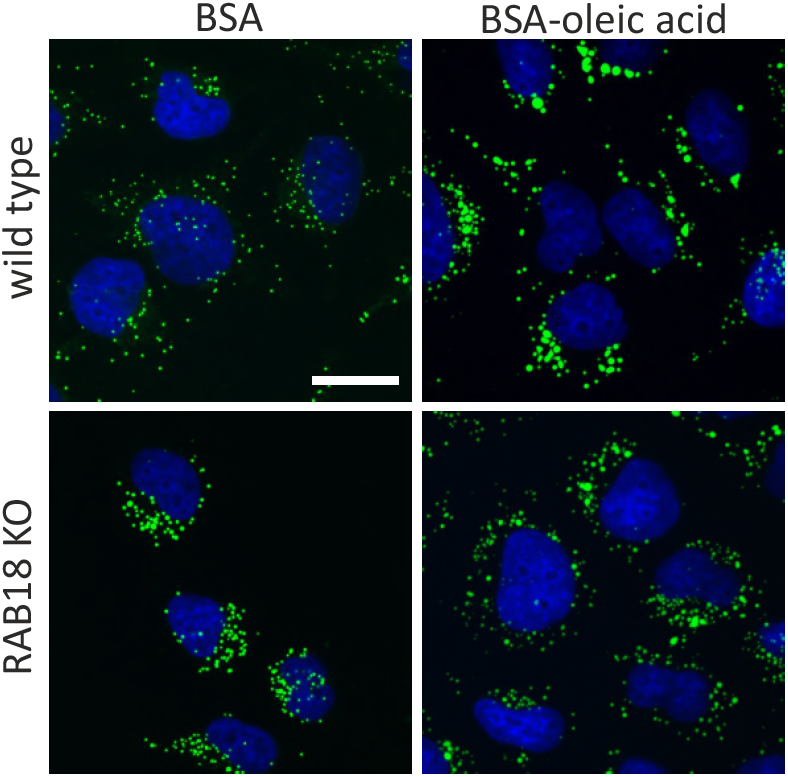
Buildup of LDs upon oleic acid treatment. Representative confocal fluorescence images of wild type and RAB18 KO cells after treatment with BSA-conjugated oleic acid or BSA alone (control). LDs are stained via BODIPY 493/503 (green) and cell nuclei by DAPI. Scale bar: 25 µm.

**Figure S3.**
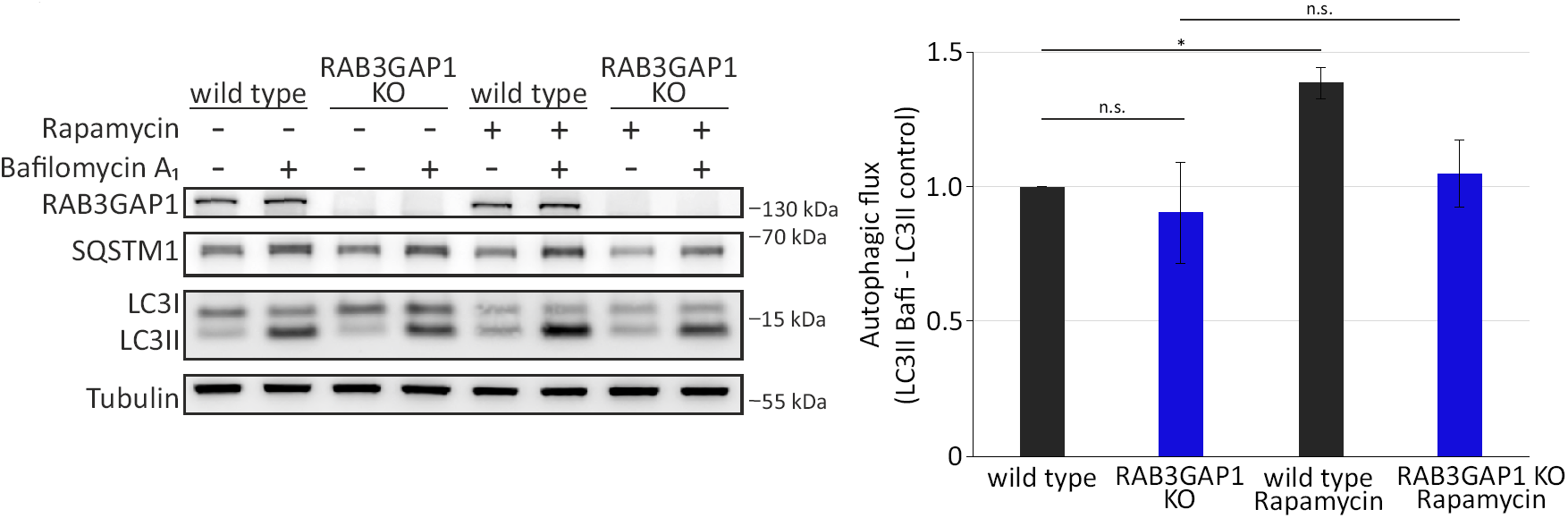
Autophagic activity in RAB3GAP1 KO cells. Immunoblot analyses of autophagic activity in wild type and RAB3GAP1 KO cells under basal and induced autophagy conditions (Rapamycin treatment). Cells were treated with DMSO (control) or Bafilomycin A1 for 2 h to allow LC3II flux evaluation. Statistics are depicted as mean +/- SD. n = 4, *t*-test, *P < 0.05.

**Figure S4.**
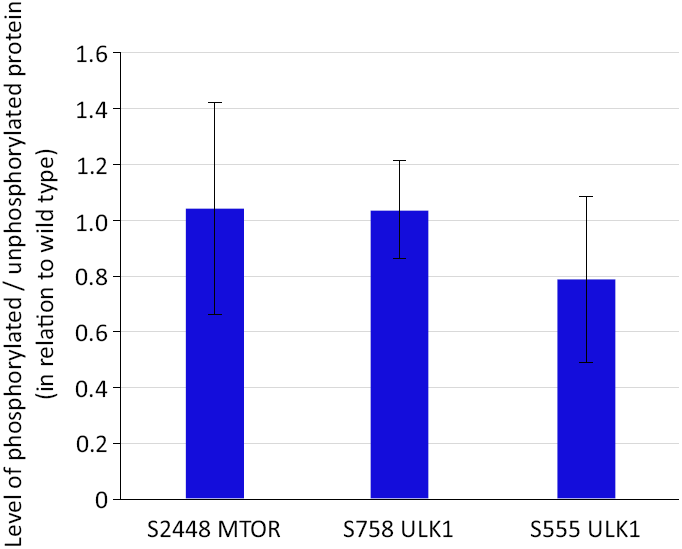
Statistical analyses of Fig. 2A. Statistics are depicted as mean +/- SD. n = 3, *t*-test. No significant differences were evaluated comparing S2448 MTOR / MTOR, S758 ULK1/ ULK1 and S555 ULK1 / ULK1 levels in wild type and RAB18 KO cells.

**Figure S5.**
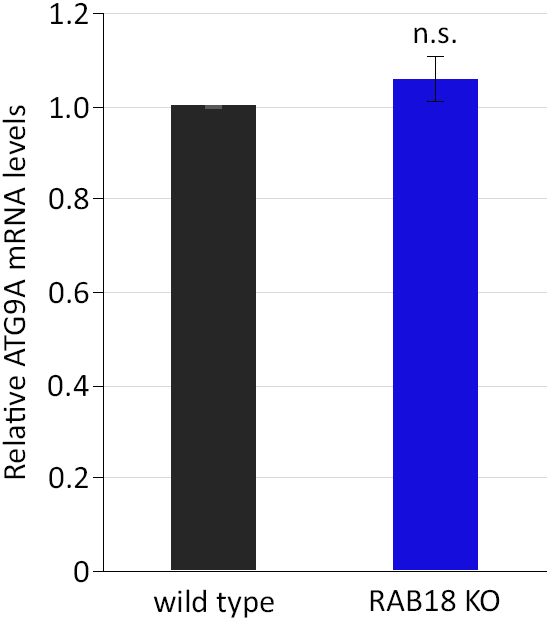
ATG9A mRNA levels are not altered in RAB18 KO cells. qPCR analyses revealed no alterations in relative ATG9A mRNA levels in comparison to wild type cells.

**Table S1.**
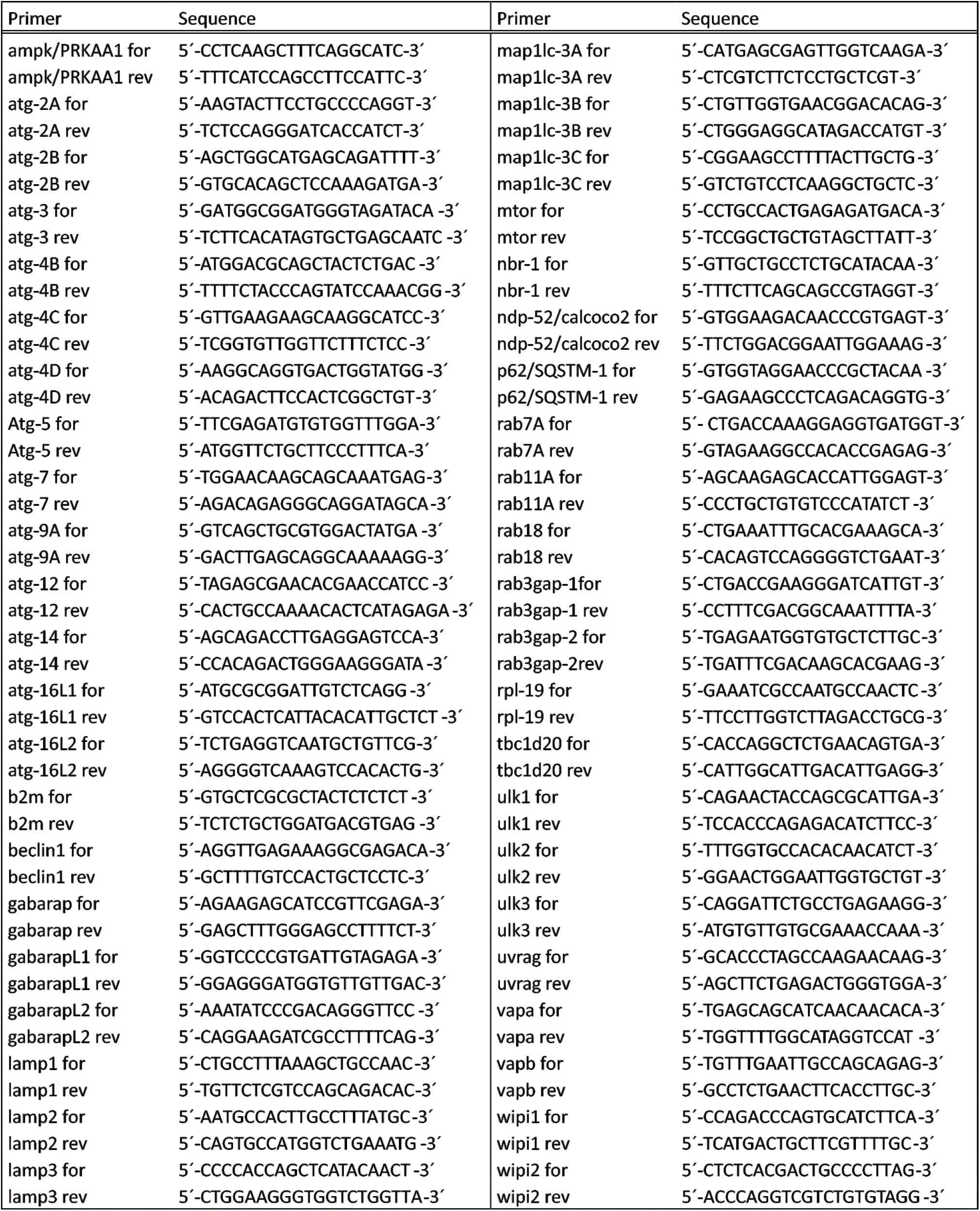
Target genes and oligo sequences of qPCR array

